# Identification of a Novel Spliced Variant of ADAMTS13 in Human Mesenchymal Stem Cells

**DOI:** 10.1101/2025.05.28.656520

**Authors:** Srishti Dutta Gupta, Nayanika Pramanik, Malancha Ta

**Author notes:** Corresponding address: Malancha Ta, PhD, Department of Biological Sciences, Indian Institute of Science Education and Research (IISER)-Kolkata, Mohanpur Campus, Dist: Nadia, West Bengal-741246, India. Extn 1217. <u>SDG</u>: Conception and design, Collection and/or assembly of data, Data analysis and interpretation, Manuscript writing; <u>NP</u>: Collection and/or assembly of data, Data analysis and interpretation; <u>MT</u>: Conception and design, Financial support, Administrative support, Data analysis and interpretation, Manuscript writing, Final approval of manuscript.

## Abstract

ADAMTS13 is a matrix metalloproteinase that cleaves von Willebrand factor (vWF) into small multimers. Several truncated forms of ADAMTS13 can be detected in plasma, generated due to alternative splicing or proteolysis by serine proteases. In this study, for the first time, we report an alternatively spliced variant of ADAMTS13 that is expressed in human mesenchymal stem cells (MSCs), derived from both the Wharton’s Jelly (WJ) of umbilical cords and the decidua-basalis layer of the placenta (PL). Our results demonstrated that the variant contained the signal peptide (Sp), propeptide (P), metalloprotease (Mp), disintegrin-like (Dis), thrombospondin type-1 repeat 1 (Tsp1-1) and cysteine-rich (Cys) domains, while the spacer (Spc) and CUB domains were absent. The Tsp1-2-8 domain was partially present. Additionally, this variant possessed partial vWF cleavage activity compared to full-length ADAMTS13. Thus, we identified a novel isoform of ADAMTS13, generated due to exon skipping in human MSCs, which retained partial activity towards vWF cleavage.

## 1. INTRODUCTION

ADAMTS13, a disintegrin and metalloproteinase with thrombospondin type-1 motif, member 13, located on the human chromosome 9q34, is a member of the zinc metalloproteinase ADAMTS family of genes (1). It has proteolytic activity that cleaves von Willebrand factor (vWF) to establish an even distribution of small multimers that is critical in regulating vascular homeostasis. A deficiency in the activity of ADAMTS13 is known to trigger a clotting disorder called thrombotic thrombocytopenic purpura (TTP) (1,2). The *ADAMTS13* gene spans 29 exons and encodes a protein with 1427 amino acids. The protein, from its N-terminus, includes multiple domains like a signal peptide (Sp), propeptide (P), metalloprotease (Mp), a disintegrin-like (Dis), a thrombospondin type-1 repeat 1 (Tsp1-1), a cysteine-rich (Cys), a spacer (Spc), seven additional thrombospondin type-1 repeats (Tsp1-2-8) and two CUB domains (3).

ADAMTS13 enables the proteolysis of the Tyr1605-Met1606 scissile bond in the A2 domain of vWF and causes its cleavage. The Mp domain of ADAMTS13, which contains the zinc binding residues, plays a crucial role in regulating this proteolysis (4). However, studies have reported that the proteolytic activity of ADAMTS13 is not just limited to its Mp domain. It has been shown that the Dis, Tsp1-1, Cys and Spc domains are also required for interacting with and cleaving vWF (5,6). In fact, the Dis domain serves to position the vWF A2 domain in the active site of ADAMTS13 to enable the proteolytic cleavage of the scissile bond (7,8). Similarly, the Spc domain is essential for recognizing and binding to vWF (9,10). Following the Spc domain, the C-terminus of ADAMTS13, comprising the Tsp1-2-8 and the two CUB domains, induces conformational changes in ADAMTS13 which help to expose the active sites (11). These are not directly involved in regulating the proteolytic activity, as corroborated by findings involving ADAMTS13 constructs truncated after the Spc domain that retain vWF cleavage activity in vitro (9,12).

Alternative splicing is an essential post-transcriptional modification leading to the generation of different types of mature mRNAs with varied structures and functions. Various mechanisms have been identified, with exon skipping being the most notable one (13). Alternative splicing is a common feature in the ADAMTS family of genes (14). Some alternatively spliced transcripts for ADAMTS13 were characterized as having different abilities to interact with the extracellular matrix or vWF. A previous study reported four probable isoforms of ADAMTS13 in the liver, resulting from two alternative splice sites, one at the Dis region and another at the COOH-terminal region (15). Similarly, another study identified seven possible alternatively spliced variants for ADAMTS13 in different tissues like liver, brain and prostate that were truncated after the Mp, Spc, or first CUB domain (16). Yet another report in hepatic stellate cells revealed the presence of an ADAMTS13 isoform, which retained the 25^th^ intron, post-splicing. Though this isoform retained the vWF cleaving activity, it was incapable of maintaining homeostasis in circulating blood (17). A recent report studying single-nucleotide variants in ADAMTS13 identified the existence of seven probable isoforms generated due to exon skipping and intron retention (3).

ADAMTS13 is mainly expressed in endothelial cells, podocytes, microglial cells and hepatic stellate cells (2). We detected the expression of ADAMTS13 in mesenchymal stem cells (MSCs) derived from birth-associated tissues like Wharton’s Jelly (WJ) of human umbilical cords and human placenta (PL). A recent finding from our group demonstrated that the ADAMTS13 expressed in WJ-MSCs was of__∼__50 kDa size and not 150–190 kDa as previously reported (18). We hypothesized that this could be due to an alternatively spliced variant of ADAMTS13 in WJ-MSCs.

Here, we identified a novel alternatively spliced variant of ADAMTS13 in WJ and PL-MSCs. Our results depicted that this variant contained exons 1 to 14, 20, 24 and 25. The exons 15 to 19, 21 to 23 and 26 to 29 were skipped in this variant, as demonstrated by exon-based PCRs. Thus, it could be concluded that the Mp, Dis, Tsp1-1 and Cys domains were present in this variant, while the Spc and CUB domains were absent. The Tsp1-2-8 domain was partially present. The presence or absence of the exons was validated by Sanger sequencing. Additionally, this variant was shown to retain vWF cleaving property by an in vitro fluorescence-based assay, thereby confirming its functional ability. To the best of our knowledge, this is the first report that suggests the existence of an alternatively spliced functional variant of ADAMTS13 in human MSCs.

## 2. MATERIALS AND METHODS

### 2.1 Cell culture

Human umbilical cord (n∼5) and placenta samples (n∼3) were collected after full-term births by caesarean delivery, with prior consent from the donor, following the guidelines laid down by the Institutional Ethics Committee (IEC) at IISER Kolkata, India. All methods were carried out according to the guidelines of the Declaration of Helsinki. MSCs were isolated from the perivascular region of the umbilical cords (WJ-MSCs) and the decidua-basalis layer of the full-term placenta (PL-MSCs) by explant culture method, as described previously (19,20). WJ and PL-MSCs were cultured in complete medium consisting of KnockOut^TM^ DMEM, supplemented with 10% MSC-qualified fetal bovine serum (FBS), 2 mM L-Glutamine or 1X GlutaMAX^TM^ and 1X Penicillin-Streptomycin (all from Thermo Fisher Scientific, USA). After 24 hrs, the medium was replaced with KnockOut^TM^ DMEM, but without any FBS, and cultured for the next 48 hrs. Meanwhile, the control cells were maintained in the complete medium for a total of 72 hrs.

### 2.2 RNA isolation and cDNA synthesis

Total RNA was isolated using RNAiso Plus (Takara, Japan) as per the manufacturer’s protocol, and the RNA yield was quantified using Nanodrop 2000 spectrophotometer (Thermo Fisher Scientific). Post RNA isolation, DNAse I treatment (Thermo Fisher Scientific) was performed according to the manufacturer’s protocol. cDNA synthesis was carried out using Verso cDNA synthesis kit (Thermo Fisher Scientific).

### 2.3 Semi-quantitative reverse-transcription polymerase chain reaction (RT-PCR)

Primers were designed for all the individual domains and exons of ADAMTS13 (listed in Table 1) and semi-quantitative RT-PCR was performed using both WJ and PL-MSC cDNAs. The PCR products were run on agarose gel. Human umbilical vein endothelial cell (HUVEC) cDNA, known to express full-length ADAMTS13, was used as a positive control. Next, the regions between exons 1 to 4, 4 to 9, 9 to 14, 14 to 20 and 20 to 25 were amplified and the specific bands were excised out.

**Table 1:**
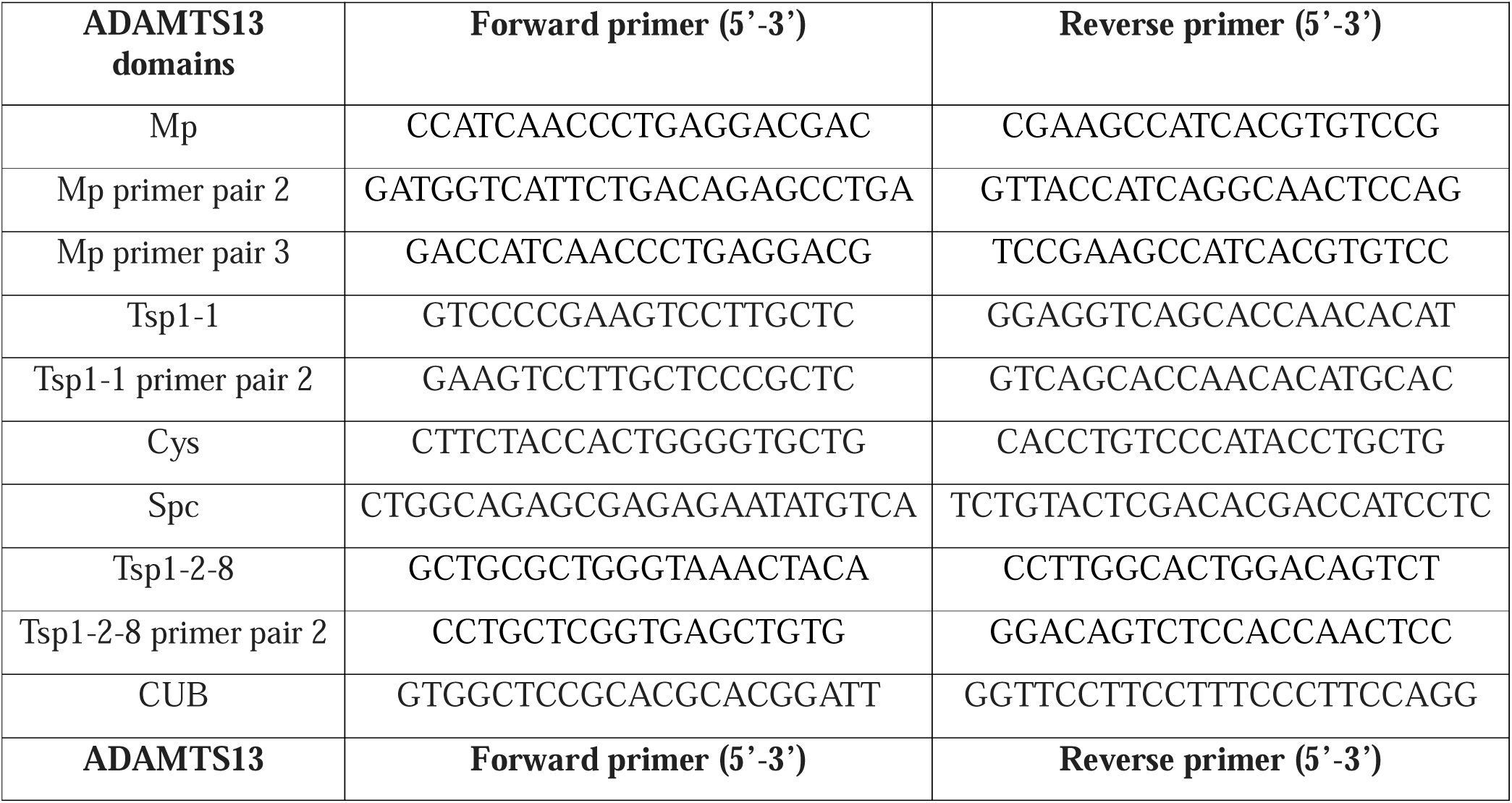
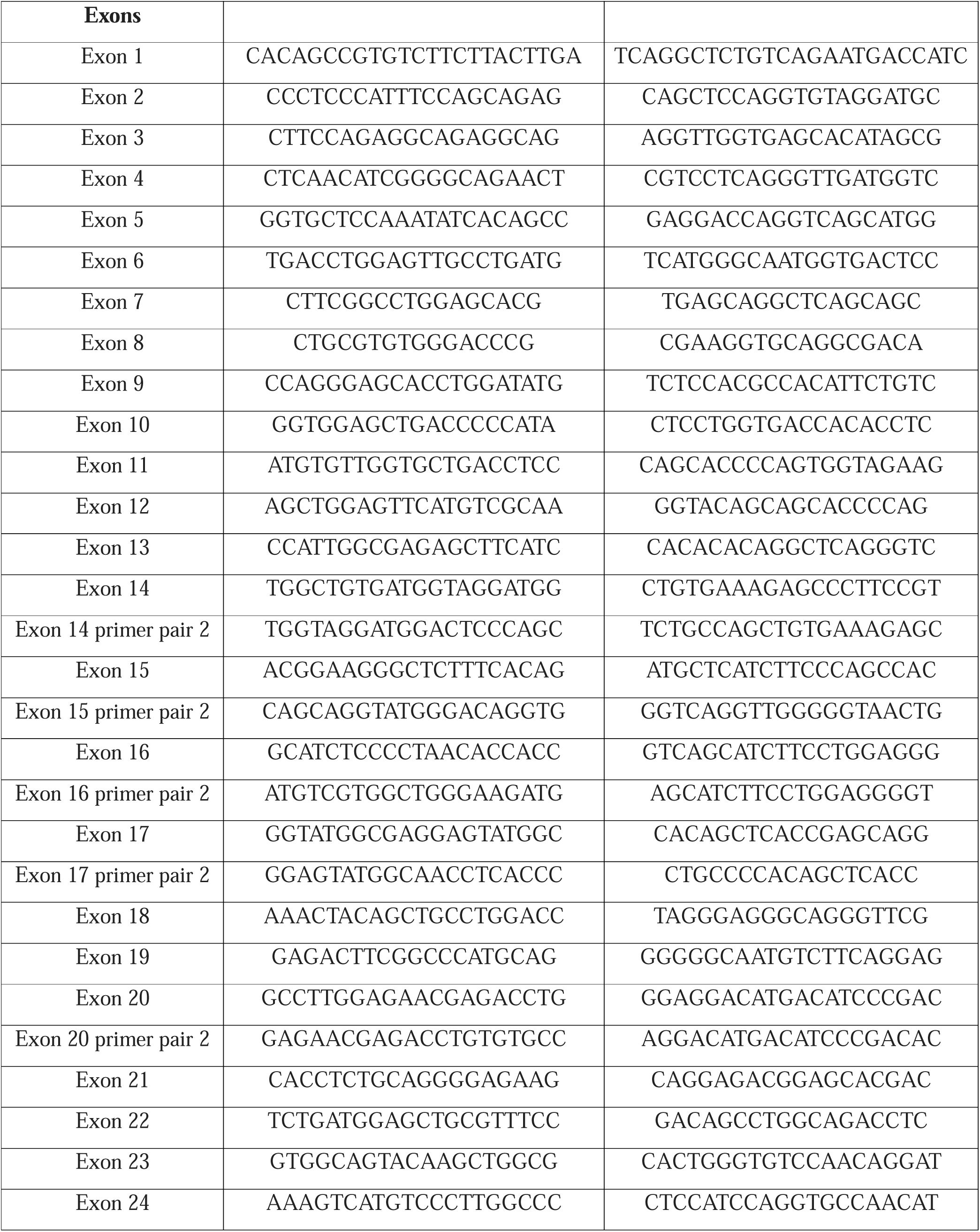
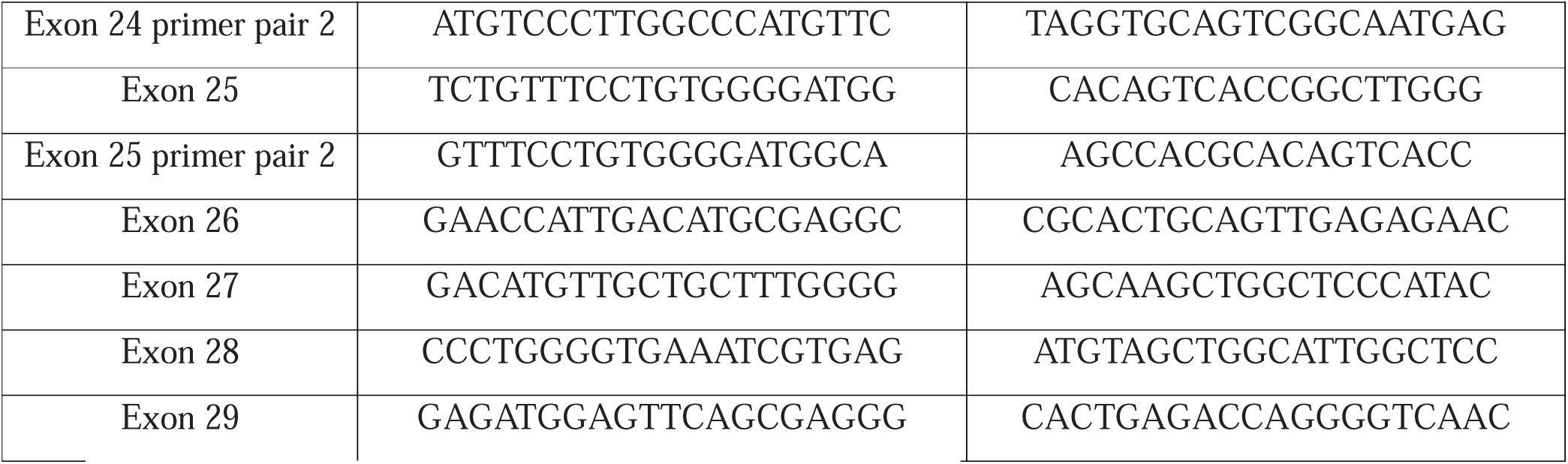
Primer sequences used for semi-quantitative RT-PCR.

### 2.4 DNA Gel elution

Gel elution of the PCR products was carried out using the QIAquick Gel Extraction Kit (QiaGen, Germany), using the manufacturer’s protocol.

### 2.5 Sanger sequence analysis

Sequences of the eluted PCR products were determined by Sanger sequencing, using the 3500 Genetic Analyzer (Applied Biosystems, USA) machine. The reads were analyzed, and sequence alignment was performed using the NCBI Basic Local Alignment Search Tool (BLAST) software.

### 2.6 Analysis of ADAMTS13 activity

ADAMTS13 activity in WJ and PL-MSC lysates was assessed using the fluorimetric SensoLyte^®^ 520 ADAMTS13 Activity Assay Kit (AnaSpec, USA), as per the manufacturer’s protocol.

### 2.7 Statistical analysis

All data are presented as mean__±__standard error of the mean (SEM). Data analysis and graphical representations were performed using GraphPad Prism 8 software (GraphPad, USA). Statistical comparisons were assessed using column statistics (one-sample t-test). Significance was accepted at *p*__<__0.05.

## 3. RESULTS AND DISCUSSION

### 3.1 Identification of the domains and exons of ADAMTS13 expressed in WJ and PL-MSCs

To identify the domains present in ADAMTS13 that is expressed by WJ and PL-MSCs, semi-quantitative PCR was performed using different domain-based exon-spanning primers. Primer pairs were designed to amplify the Mp, Tsp1-1, Cys, Spc, Tsp1-2-8 and CUB domains of ADAMTS13. Interestingly, bands were observed only for the Mp, Tsp1-1 and Cys domains (**Fig. 1A**). Amplification of the Spc, Tsp1-2-8 and CUB domains did not yield any bands (**Fig. 1A**). HUVECs, known to express the full-length ADAMTS13, showed the presence of all the domains (**Fig. 1A**). More than one primer pair was used to confirm the presence or absence of expression of each of the following Mp, Tsp1-1 and Tsp1-2-8 domains and similar results were obtained (**Supplementary** Fig. 1A), indicating that only these domains were present in the ADAMTS13 expressed by WJ and PL-MSCs. cDNAs prepared from WJ and PL-MSCs, cultured under both control and serum-deprivation conditions, were used as a part of this study and similar results were observed.

**Fig. 1:**
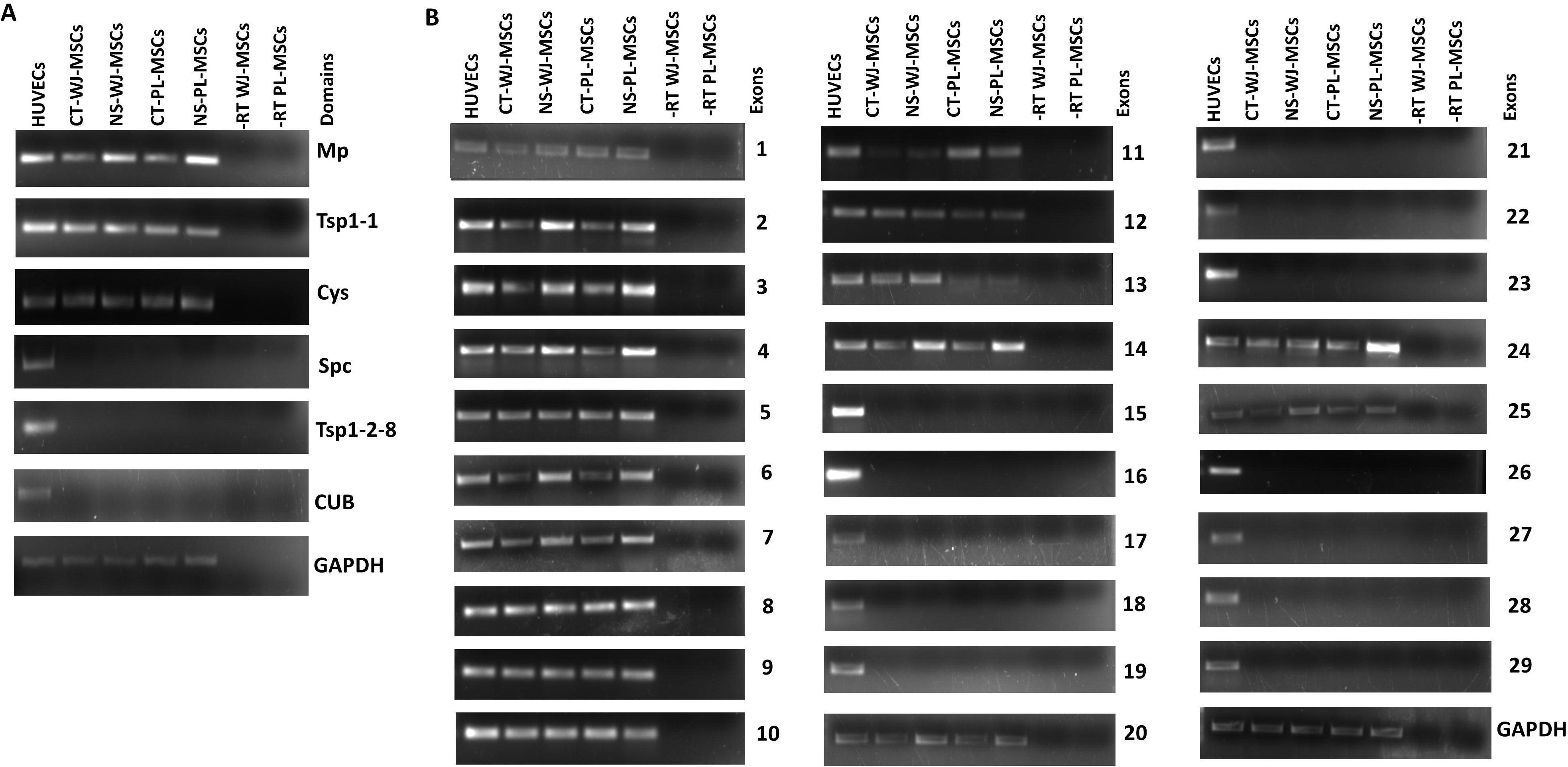
Identification of the domains and exons of ADAMTS13 expressed in WJ and PL-MSCs. Semi-quantitative RT-PCRs were performed to detect the presence or absence of the various domains/exons of ADAMTS13 in WJ and PL-MSCs. (**A**) Exon-spanning primer pairs were designed to amplify the Mp, Tsp1-1, Cys, Spc, Tsp1-2-8 and the CUB domains of ADAMTS13 and representative gel images are shown (n = 2). (**B**) Primers were designed to amplify each of the 29 individual exons of ADAMTS13 and representative gel images are shown (n = 2). GAPDH was used as the endogenous control. HUVEC cDNA, known to express full-length ADAMTS13, was used as a positive control. A set of WJ and PL-MSC cDNAs prepared without the reverse transcriptase enzyme was used as the negative control (-RT) for our experiments. Serum-deprivation condition has been denoted as no serum (NS) and control as CT in the figure.

For further validation, PCRs were carried out for each of the 29 individual exons of ADAMTS13. Specific bands were obtained upon amplifying the exon numbers 1, 2, 3, 4, 5, 6, 7, 8, 9, 10, 11,12, 13, 14, 20, 24 and 25 in WJ and PL-MSCs cultured under both control and serum-deprivation conditions. However, the exons 15, 16, 17, 18, 19, 21, 22, 23, 26, 27, 28 and 29 were found to be missing (**Fig. 1B**). Again, HUVECs showed a band for PCR amplification for each of the 29 exons (**Fig. 1B**). More than one primer pair was used to verify the presence or absence of some of the exons (**Supplementary** Fig. 1B). These observations corroborated the presence of the domains and their corresponding exons as shown in the Table 2.

**Table 2:** The various domains of ADAMTS13 and their corresponding exons that are present or absent in the ADAMTS13 variant observed in WJ and PL-MSCs.

| Domains | Corresponding exons in the variant |  |
| --- | --- | --- |
|  | Present | Absent |
| Sp | 1 | - |
| P | 2 | - |
| Mp | 3-8 | - |
| Dis | 8-10 | - |
| Tsp1-1 | 10-12 | - |
| Cys | 12-14 | - |
| Spc | 14 | 15-17 |
| Tsp1-2-8 | 20, 24, 25 | 17-19, 21-23 |
| CUB | - | 26-29 |

These observations suggested the existence of an alternatively spliced variant of ADAMTS13, generated by exon skipping, in WJ and PL-MSCs.

### 3.2 Confirmation of the exons present in the alternatively spliced variant by Sanger sequencing

As WJ and PL-MSCs, cultured under both control and serum-deprivation conditions, demonstrated the expression of the same domains/exons of ADAMTS13, the next part of the work was performed only with the control WJ-MSC cDNA. To confirm the presence of the exons by Sanger sequencing, PCR amplification was next performed for exons 1-4, using the forward primer specific for exon 1 and the reverse primer specific for exon 4. Similarly, PCR products were generated for exons 4-9, 9-14, 14-20 and 20-25, gel purified and sequenced.

The sequencing data, when aligned against the full-length ADAMTS13 sequence, confirmed the synchronous presence of exons 1-4, 4-9 and 9-14 without any exons missing in between (**Fig. 2, Supplementary** Figure 2-4). However, the sequencing data for exons 14-20, when aligned against the full-length ADAMTS13 sequence, showed specific pairing at two regions corresponding to exon 14 and exon 20 only, and the exons 15-19 were missing in between (**Fig. 2, Supplementary** Figure 5). Similarly, the sequencing data for exons 20-25, when aligned against the full-length ADAMTS13 sequence, showed specific pairing corresponding to exons 20, 24 and 25 while the exons 21-23 were missing in between (**Fig. 2, Supplementary** Figure 6). In accordance with the exon-based PCR data, the sequencing data confirmed the skipping of exons 15-19 and 21-23 in the alternatively spliced variant of ADAMTS13.

**Fig. 2:**
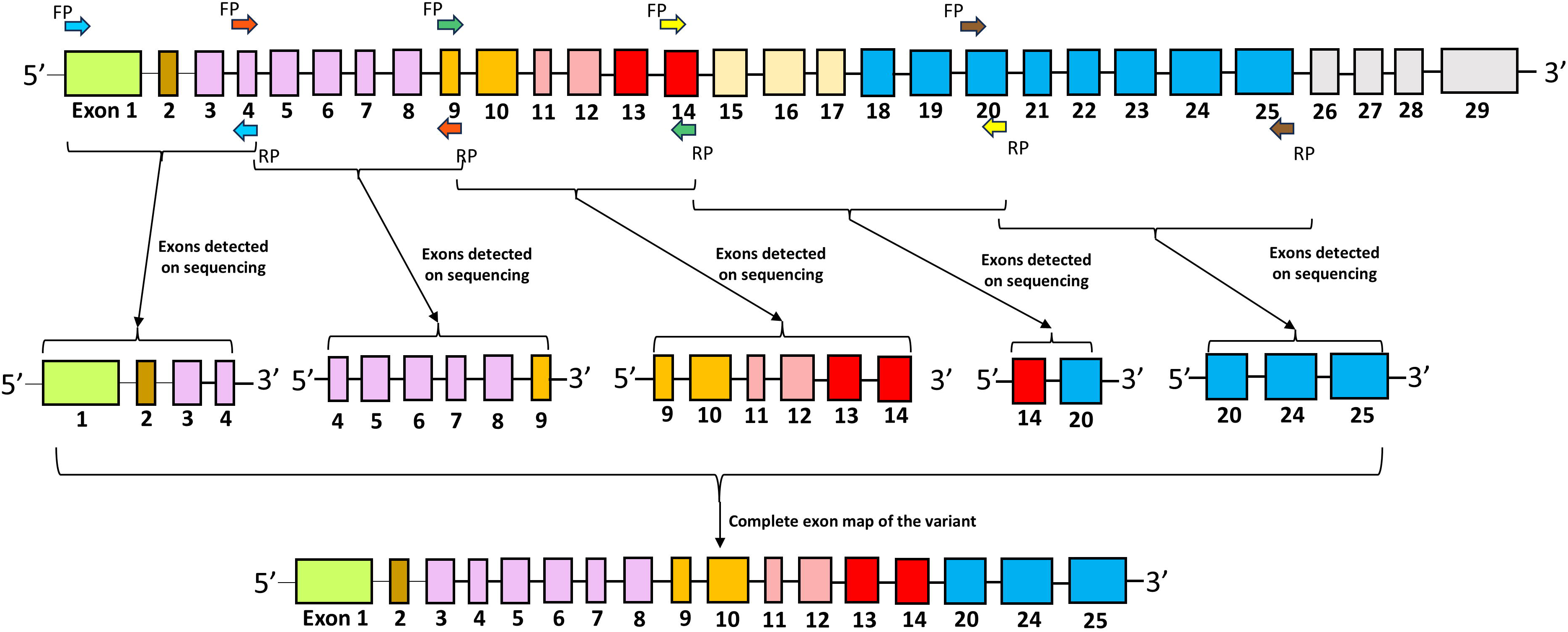
Presenting the PCR amplification map for Sanger sequencing. PCR was performed to amplify ADAMTS13 between exons 1-4, 4-9, 9-14, 14-20 and 20-25, using their respective forward and reverse primers, as shown in the figure. Arrows of the same colour have been used to denote the forward and reverse primer pairs used in a particular amplification. The amplified products, so obtained, were gel-purified and sequenced to confirm the exons present or absent in the variant. The forward and reverse primers have been denoted as FP and RP, respectively, in the figure.

### 3.3 Assessment of the vWF cleavage activity of the alternatively spliced variant of ADAMTS13

To evaluate the functional activity of the spliced variant, an internally quenched vWF73 FRET peptide was employed which would emit a fluorescence signal upon cleavage by ADAMTS13. In this in vitro assay, the vWF73 FRET peptide was added to WJ or PL-MSC lysates and their endogenous vWF cleavage activity was measured by recording the fluorescence emission signal in relative fluorescence units (RFUs). Recombinant human full-length ADAMTS13 (rhADAMTS13), which has vWF cleavage activity, was used as a positive control in this experiment. The WJ and PL-MSC lysates emitted fluorescence signals of intensities 5133 ± 151.9 RFUs and 3762 ± 997.3 RFUs, respectively (**Fig. 3A**). Meanwhile, rhADAMTS13 emitted a fluorescence intensity of 10457 ± 1200 RFUs (**Fig. 3A**). These results indicated that the alternatively spliced variant in WJ and PL-MSCs possessed partial activity, ∼ 50% and 36%, respectively, relative to rhADAMTS13 (**Fig. 3B**). This reduction in activity could possibly be due to the absence of the Spc domain in this variant, as depicted in our previous results. Lending support to our observation, a previous study had reported that an ADAMTS13 construct containing the Mp, Dis, Tsp1-1 and Cys domains possessed only half the relative activity of full-length ADAMTS13, while addition of the Spc domain to the construct restored the complete activity (21). Another report suggested that deleting the Spc domain from ADAMTS13 reduced the rate of vWF cleavage by ∼ 20-fold (22).

**Fig. 3:**
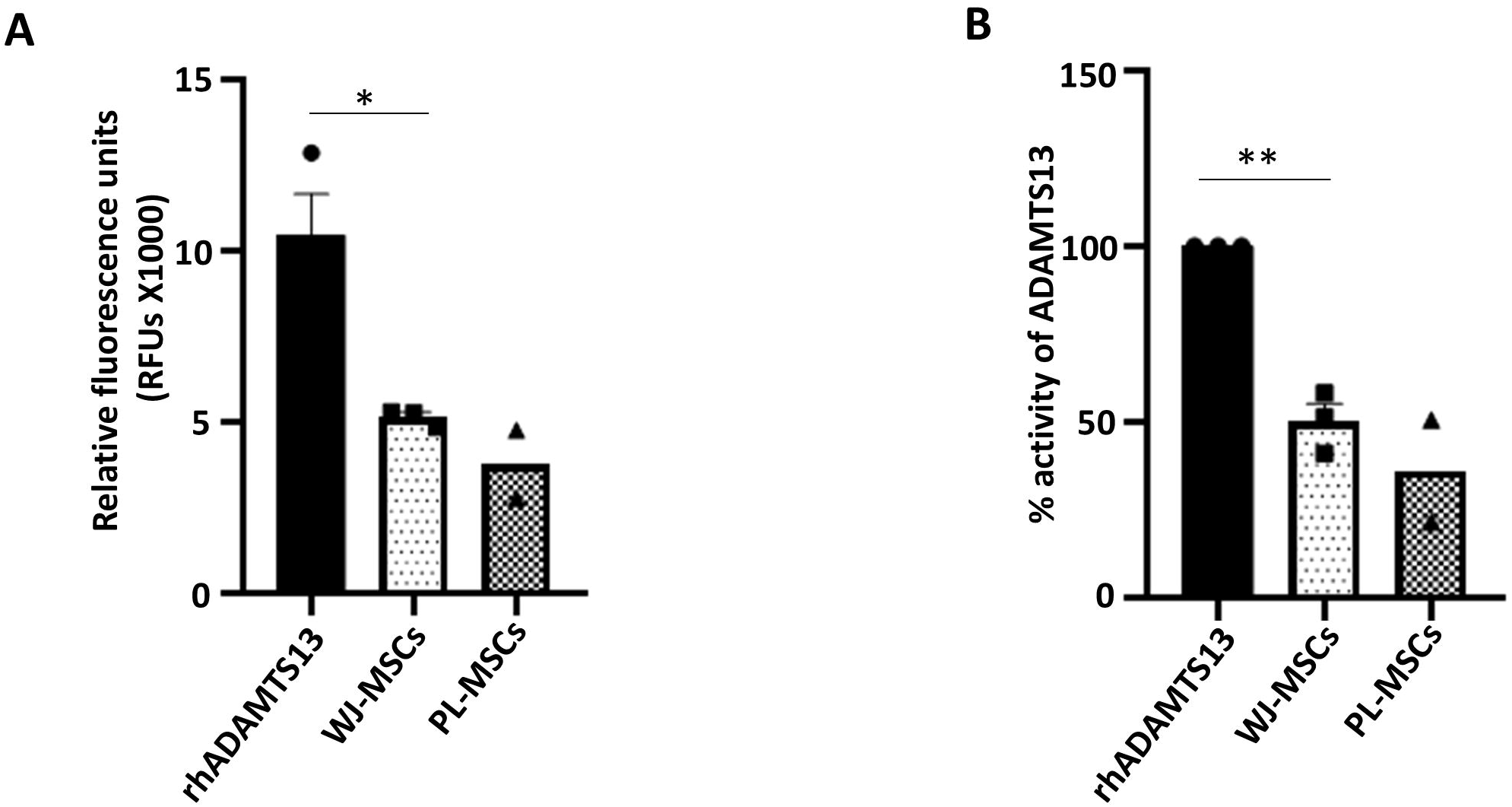
Assessment of the vWF cleavage activity of the alternatively spliced variant of ADAMTS13. (**A**) To measure the endogenous vWF cleavage activity of WJ and PL-MSC lysates, the intensity of their fluorescence emission signal was recorded in RFUs and plotted (n ≥ 2). (**B**) The percentage of vWF cleavage activity of the alternatively spliced variant, relative to recombinant human full-length ADAMTS13 (represented as rhADAMTS13), was also calculated and plotted (n ≥ 2). Each bar represents mean[±[SEM (* *p*[<[0.05, ** *p*[<[0.01).

Overall, our findings highlight the presence of a novel alternatively spliced variant of ADAMTS13 in MSCs derived from birth-associated tissues. However, further in-depth studies would be needed to elucidate the substrate specificity and mechanism of proteolytic processing in this variant.

## ACKNOWLEDGEMENTS

The work was funded by DBT (Project ID-BT/PR45601/MED/31/460/2022) and IISER Kolkata. We thank UGC, India for the fellowship of Ms Srishti Dutta Gupta. We are grateful to Dr. Jayanta Chatterjee, Astha, Kalyani for generously providing the umbilical cord and placenta samples. We are thankful to Mr. Tamal Ghosh and Mr. Somnath Halder for technical assistance with flow cytometry and Sanger sequencing. We thank Ms. Ankita Sen and Mr. Pritam Banerjee for their help with tissue collection and MSC isolation.

