## Supplemental file for "Identification of a Novel Spliced Variant of ADAMTS13 in Human Mesenchymal Stem Cells"

**Supplementary data**

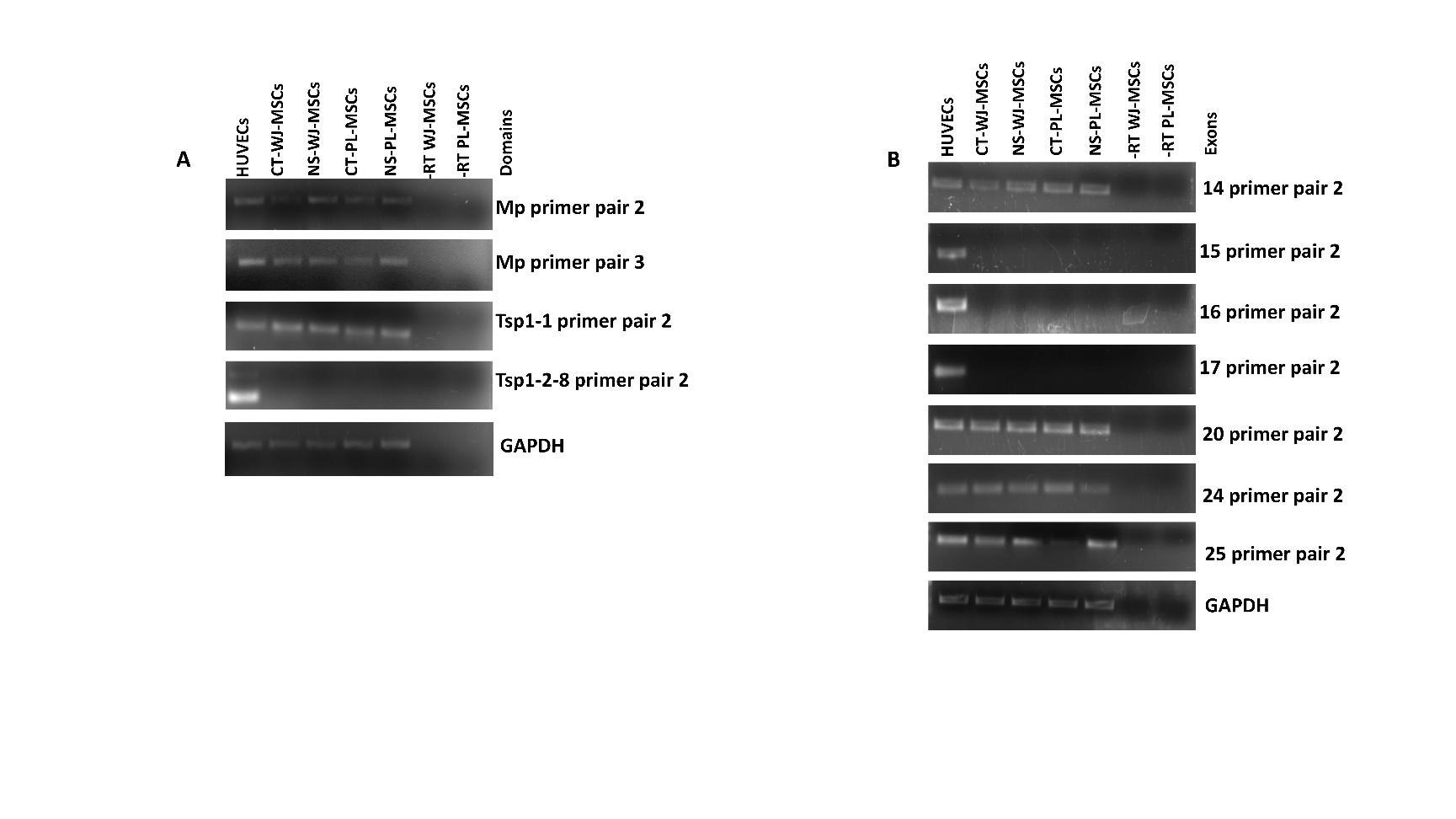

**Supplementary Figure 1:** Identification of the domains and exons of ADAMTS13 expressed in WJ and PL-MSCs. Semi-quantitative RT-PCRs were performed to detect the presence or absence of the various domains/exons of ADAMTS13 in WJ and PL-MSCs. (**A**) More than one primer pairs were designed to amplify the Mp, Tsp1-1 and Tsp1-2-8 domains of ADAMTS13 and representative gel images are shown (n = 2). (**B**) Similarly, more than one primer pairs were designed to amplify exons 14, 15, 16, 17, 20, 24 and 25 of ADAMTS13 and representative gel images are shown (n = 2). GAPDH was used as the endogenous control. HUVEC cDNA, known to express full-length ADAMTS13, was used as a positive control. A set of WJ and PL-MSC cDNAs prepared without the reverse transcriptase enzyme was used as the negative control (-RT) for our experiments. Serum-deprivation condition has been denoted as no serum (NS) and control as CT in the figure.

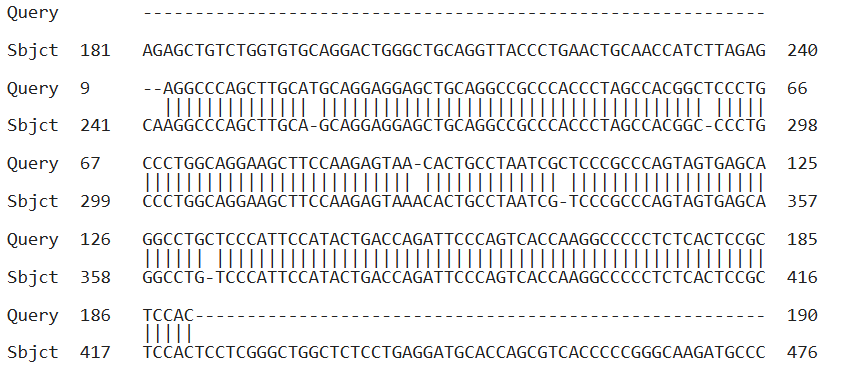

**Supplementary Figure 2:** Sequence of the PCR amplified product of ADAMTS13 between exons 1 to 4, aligned against the full-length ADAMTS13 sequence, is represented. The synchronous presence of exons 1-4 was observed without any exons missing in between. cDNA prepared from WJ-MSCs cultured under control condition was used to perform the PCR.

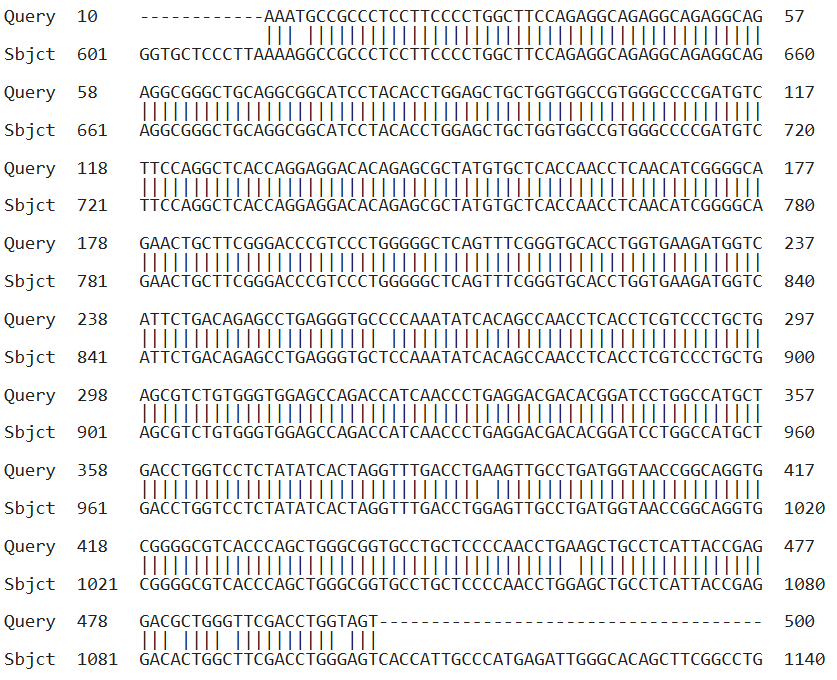

**Supplementary Figure 3:** Sequence of the PCR amplified product of ADAMTS13 between exons 4 to 9, aligned against the full-length ADAMTS13 sequence, is represented. The synchronous presence of exons 4-9 was observed without any exons missing in between. cDNA prepared from WJ-MSCs cultured under control condition was used to perform the PCR.

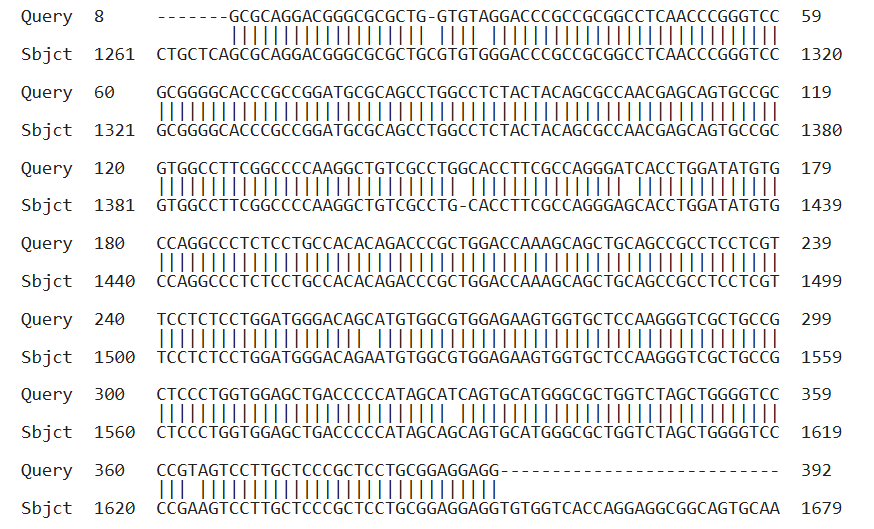

**Supplementary Figure 4:** Sequence of the PCR amplified product of ADAMTS13 between exons 9 to 14, aligned against the full-length ADAMTS13 sequence, is represented. The synchronous presence of exons 9-14 was observed without any exons missing in between. cDNA prepared from WJ-MSCs cultured under control condition was used to perform the PCR.

.

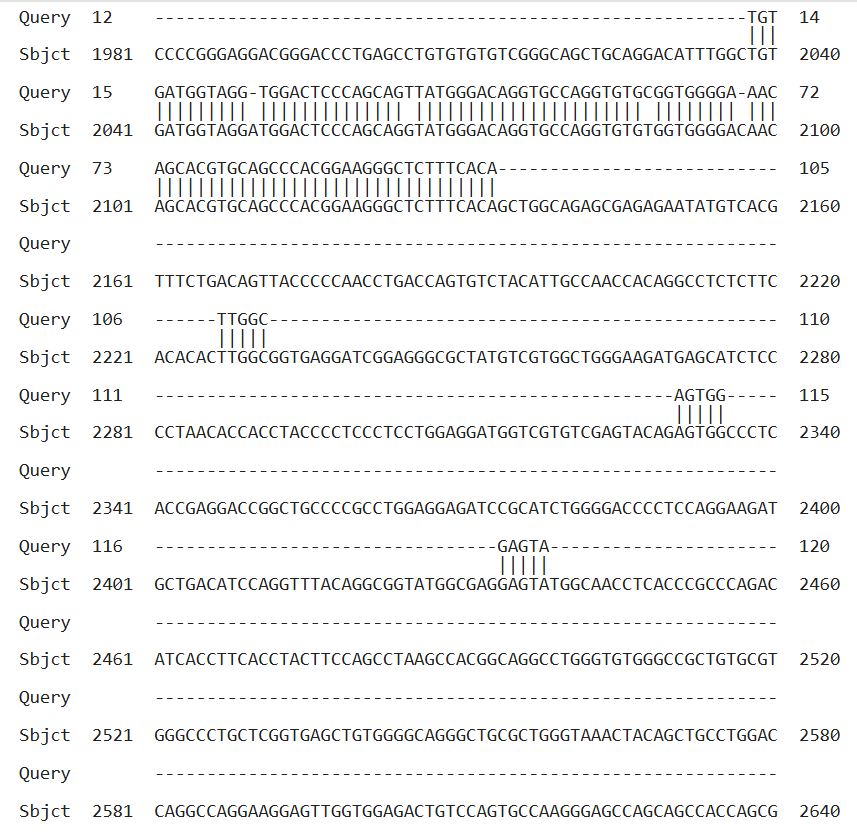

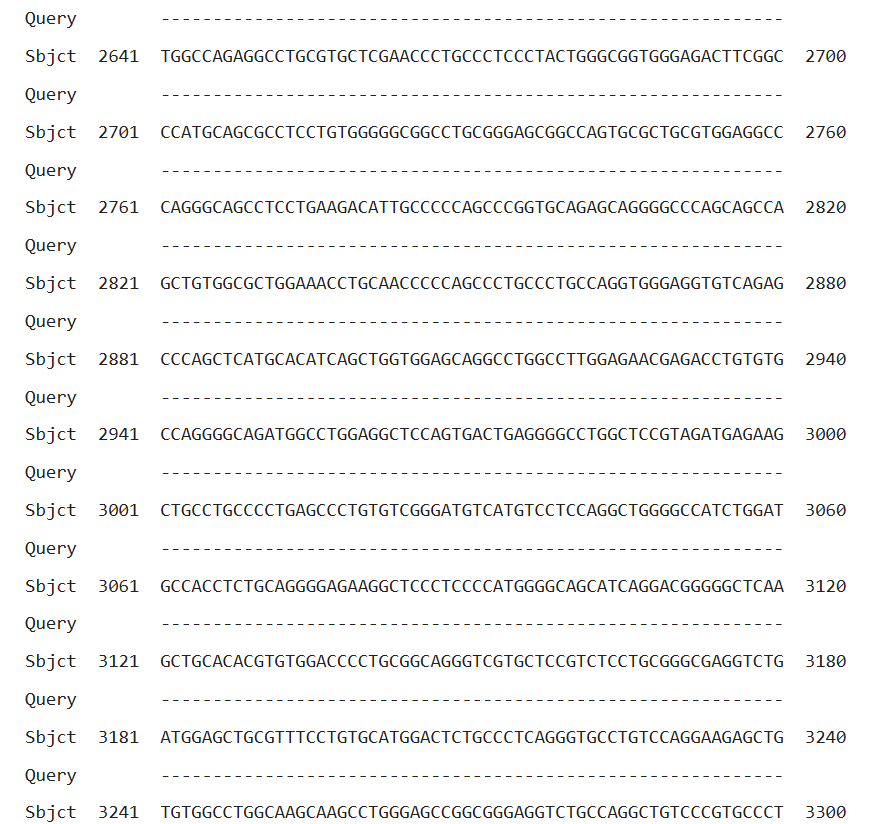

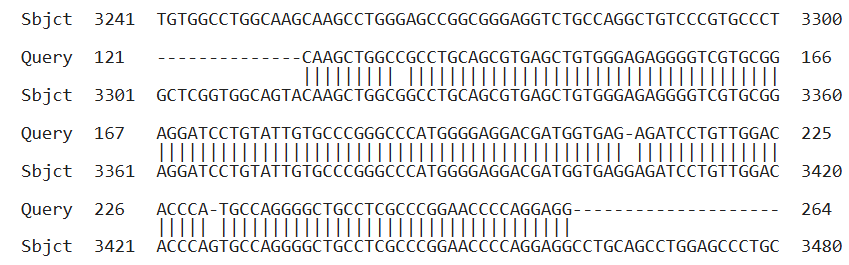

**Supplementary Figure 5:** Sequence of the PCR amplified product of ADAMTS13 between exons 14 to 20, aligned against the full-length ADAMTS13 sequence, is represented. The boxes outlined in red depict the regions of specific alignment pertaining to exons 14 and 20 only, while exons 15-19 are missing in between. cDNA prepared from WJ-MSCs cultured under control condition was used to perform the PCR.

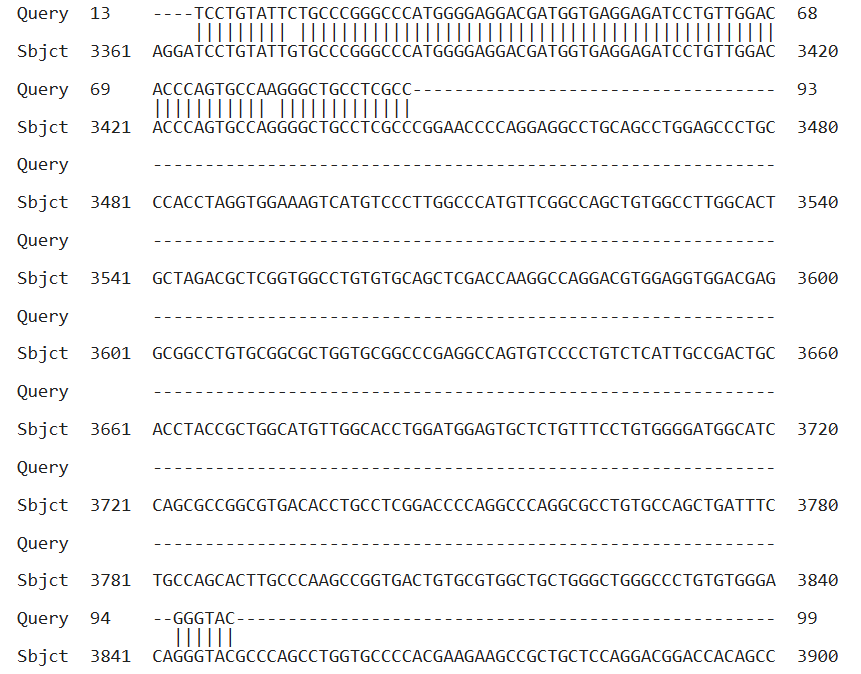

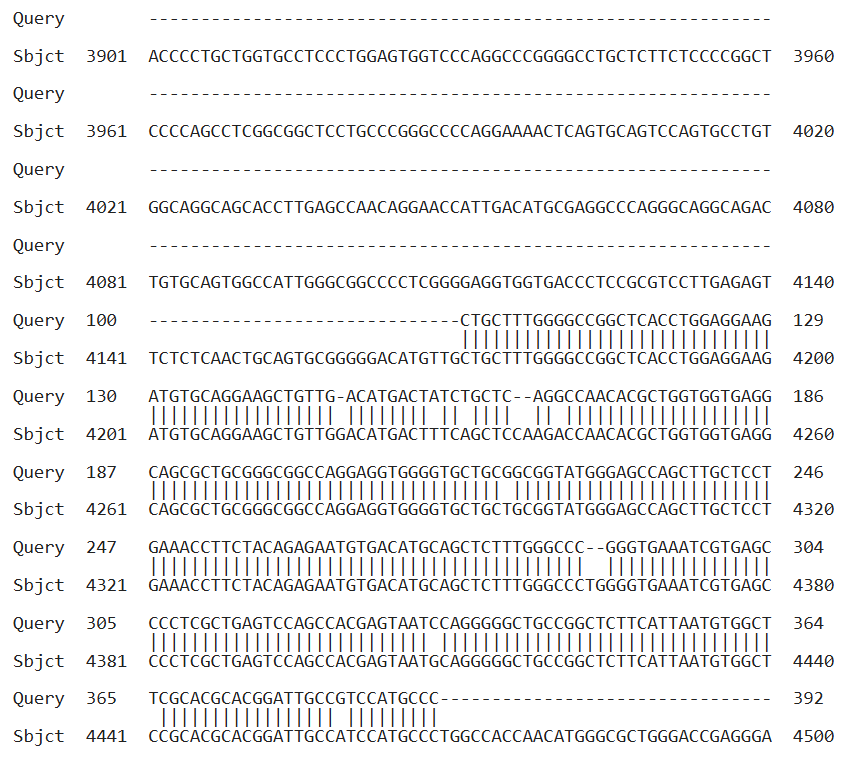

**Supplementary Figure 6:** Sequence of the PCR amplified product of ADAMTS13 between exons 20 to 25, aligned against the full-length ADAMTS13 sequence, is represented. The boxes outlined in red depict the regions of specific alignment pertaining to exons 20, 24 and 25, while exons 21-23 are missing in between. cDNA prepared from WJ-MSCs cultured under control condition was used to perform the PCR.
